# Prefusion conformation of SARS-CoV-2 receptor-binding domain favours interactions with human receptor ACE2

**DOI:** 10.1101/2021.04.22.441041

**Authors:** Nitesh Kumawat, Andrejs Tucs, Soumen Bera, Gennady N. Chuev, Marina V. Fedotova, Koji Tsuda, Sergey E. Kruchinin, Adnan Sljoka, Amit Chakraborty

## Abstract

A new coronavirus pandemic COVID-19, caused by Severe Acute Respiratory Syndrome coronavirus (SARS-CoV-2), poses a serious threat across continents, leading the World Health Organization to declare a Public Health Emergency of International Concern. In order to block the entry of the virus into human host cells, major therapeutic and vaccine design efforts are now targeting the interactions between the SARS-CoV-2 spike (S) glycoprotein and the human cellular membrane receptor angiotensin-converting enzyme, hACE2. By analyzing cryo-EM structures of SARS-CoV-2 and SARS-CoV-1, we report here that the homotrimer SARS-CoV-2 S receptor-binding domain (RBD) that binds with hACE2 has expanded in size, undergoing a large conformational change relative to SARS-CoV-1 S protein. Protomer with the up-conformational form of RBD, which binds with hACE2, exhibits higher intermolecular interactions at the RBD-ACE2 interface, with differential distributions and the inclusion of specific H-bonds in the CoV-2 complex. Further interface analysis has shown that interfacial water promotes and stabilizes the formation of CoV-2/hACE2 complex. This interaction has caused a significant structural rigidification, favoring proteolytic processing of S protein for the fusion of the viral and cellular membrane. Moreover, conformational dynamics simulations of RBD motions in SARS-CoV-2 and SARS-CoV-1 point to the role in modification in the RBD dynamics and their likely impact on infectivity.

## Introduction

Severe Acute Respiratory Syndrome coronavirus (SARS-CoV-2) is associated with the current disease outbreak COVID-19, which has spread to more than 200 countries, surpassing 170 million infections worldwide. The first coronavirus outbreak of SARS-CoV-1 occurred in 2002 and the Middle East Respiratory Syndrome coronavirus (MERS-CoV) in 2012. Unlike previous outbreaks, rapid human-to-human transmission of SARS-CoV-2 across the continents resulted in the World Health Organization (WHO) declaration of Public Health Emergency of International Concern on January 30^th^, 2020. Although there is a large heterogeneity in disease transmission among coronaviruses, SARS-CoV-2 spread across the globe much more rapidly in comparison to the previous outbreaks [1,2]. Here we identify essential structural differences that contribute to higher transmissibility of SARS-CoV-2 relative to SARS-CoV-1.

Coronavirus (CoV) entry into human host cell is mediated by the transmembrane spike (S) glycoprotein protruding from the viral surface. With the use of S protein, which is a trimeric class I fusion protein, CoV hijacks the human receptor membrane protein angiotensin-converting enzyme 2(hACE2), which is highly expressed in lungs, heart, kidneys and intestine cells [3]. The S protein protomer is made of two subunits, S1 and S2, which are responsible for receptor recognition and membrane fusion. The S1 unit, which comprises the receptor-binding domain (RBD), binds to the peptidase domain (PD) of hACE2, and contributes to stabilization of the prefusion conformational state. Receptor-binding destabilizes the prefusion state, resulting in shedding of S1 unit and priming of S2 for membrane fusion. S-protein is then cleaved further by the host proteases at S2’ site located immediately upstream of the fusion peptide, triggering the protein for membrane fusion with stable conformational modification [4]. This essential entry process is an important target for antibody mediated neutralization.

SARS-CoV-2 S protein holds 98% sequence similarity with the bat coronavirus RaTG13. A most critical variation observed in CoV-2 is an insertion “RRAR” at the furin recognition site at S1/S2 junction, while SARS-CoV-1 has single arginine at that site. Besides this insertion, ~60% residual substitutions are noted at the RBD domain [5]. Examining how such differences contribute to higher recognition capability of hACE2 receptor is important to underpin the therapeutic target that can prevent virus entry. Here, we have analysed respective protomers of four S-protein cryo-EM structures: SARS-CoV-1 (pdb id: 6CRZ) [6], SARS-COV-2 (pdb id: 6VYB) [4] (Fig.1), SARS-CoV-RBD-hACE2 complex (pdb id: 2AJF) [7], and SARS-CoV-2-RBD-hACE2 complex (pdb id: 6M17) [8]. Recent work has revealed that the S-proteins in the open state with at least one RBD in the “up” conformation, corresponds to the receptor accessible conformation that can bind to hACE2. In comparison to the CoV-RBD up state, we show that CoV-2-RBD has expanded its net surface, undergoing a large local conformational change, higher backbone RMSD, and a relatively large amplitude anti-phase-like RBD motion in the slow-motion second normal mode. Furthermore, tight interactions between CoV-2-RBD and hACE2 have caused a significant structural rigidification, triggering proteolytic processing of S protein for membrane fusion.

**Fig.1.**
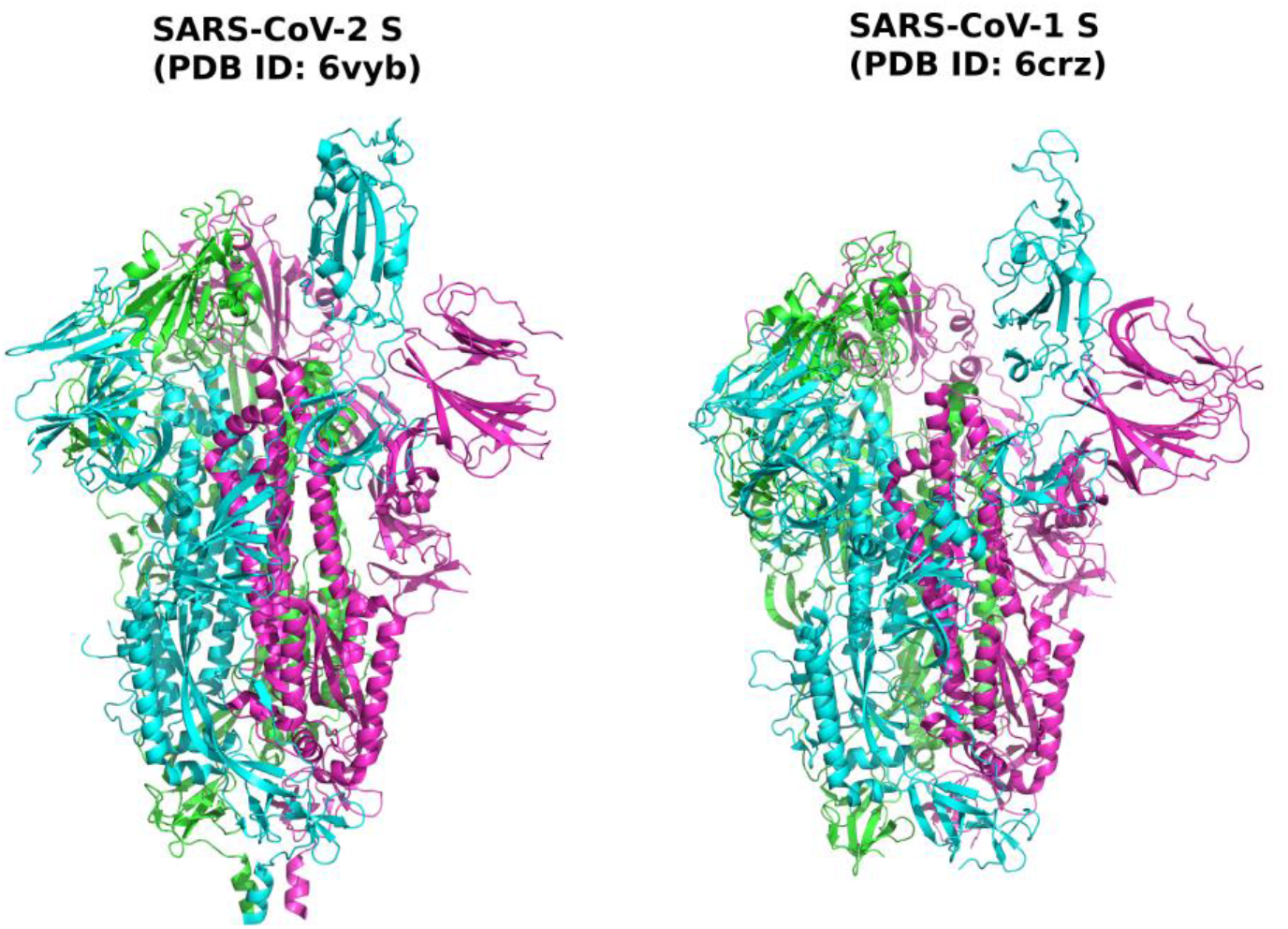
Spike (S) glycoprotein of SARS-CoV-1 and SARS-CoV-2 is a trimeric structure (indicated with three different colours), protruding from virion surface, and is responsible for the receptor recognition and membrane fusion. Here, both spike proteins have the receptor-binding domain (RBD) in the up-conformation (cyan colour) in one monomer, which can bind with the human receptor hACE2.

## Results and Discussion

### High variability of the receptor-binding domain (RBD)

Coronavirus (CoV) spike glycoprotein (S protein) protrudes from virion surface and is responsible for the receptor recognition and membrane fusion, which are essential steps for viral entry into the host cell [9]. The S1 unit of S protein contributes to receptor recognition and priming of S2 unit for membrane fusion. In order to recognize the host receptor, the receptor-binding domain (RBD) of S1 undergoes hinge-like conformational motions. In a recently reported structure of SARS-CoV-2, it was observed that human angiotensin-converting enzyme 2(hACE2) can only bind when the RBD (residues 336-518) adapts an open up-conformational state. Peptidase domain (PD) of hACE2 clashes when all the RBD domains of the homotrimer SARS-CoV-2 S are in down conformational state. To identify characteristic differences between the up-RBD of SARS-CoV-1 and CoV-2, we superimpose both the structures with and without the hACE2 (Fig.1A). Despite a number of sequence variations, structural alignments show high similarities (Fig.2A) throughout the structure. However, RBD AA residues exhibit a ~2-fold higher root mean squared deviation (RMSD) from the average level of ~2.0 Å. In contrast, bound RBD has a high RMSD variation (Fig.2B) and shows lower than the average value for many residues. Moreover, Ramachandran plot for bound-RBD-hACE2 complex displays large changes in phi and psi dihedral angles for most of the RBD residues in favourable and allowed regions of the plot (Fig.4B). These analyses indicate characteristic conformational differences in hACE2-bound and non-bound RBD between SARS-CoV-1and SARS-CoV-2.

**Fig.2.**
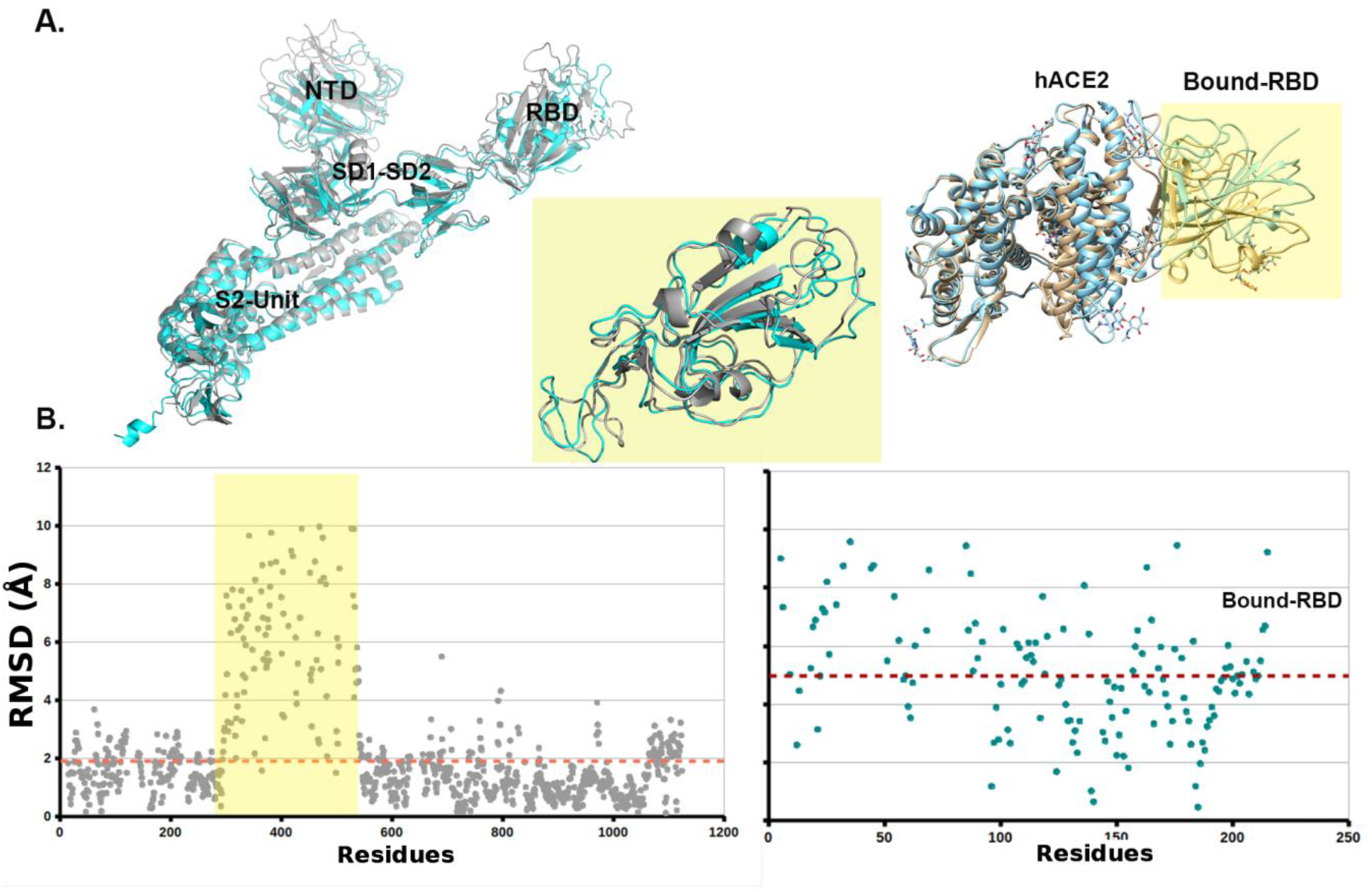
Root-mean-squared deviation (RMSD) of Cα atoms. **(A)** We used MatchMaker in UCSF-Chimera to superimpose structures. It considers similar structures to be superimposed while there are large sequence dissimilarities by producing residue pairing with the inputs of AA sequences and secondary structural elements. After superposition, it uses the usual Euclidian distance formula for calculating RMSD in angstrom unit. (B) Left side graph shows RMSD between the structures SARS-CoV-1(PDB id: 6CRZ) and CoV-2-RBD (pdb id: 6VYB) up-conformation protomer, while the right side graph shows RMSD of hACE2-bound RBD between SARS-CoV-1-RBD-hACE2 (2AJF) and SARS-CoV-2-RBD-hACE2 (6M17) complex. Yellow-shaded region shows relatively high deviations in the receptor-binding domain (RBD).

### Differential RBD motions between CoVs

Significant conformational changes are commonly reflected in the differential domain motions. To capture such differences, we have conducted normal mode analysis using the Gaussian network model (GNM) and the anisotropic-network model (ANM), utilized in DynOmics webserver and in iMODS (http://imods.chaconlab.org). This hybrid model has efficiently captured the dynamic differences between SARS-CoV-1 and SARS-CoV-2 with and without hACE2 bound. With the trimeric CoV-1 and CoV-2 macromolecular structure, we have calculated covariance matrix which is computed using the *Cα* Cartesian coordinates and the Karplus equation of collective motion in protein. This covariance matrix signifies coupling between pairs of residues (Fig.3A,B). Overall, covariance patterns are very similar for CoV-1 and CoV-2; however, there are few sharp differences in few spots of the covariance matrices.

**Fig.3.**
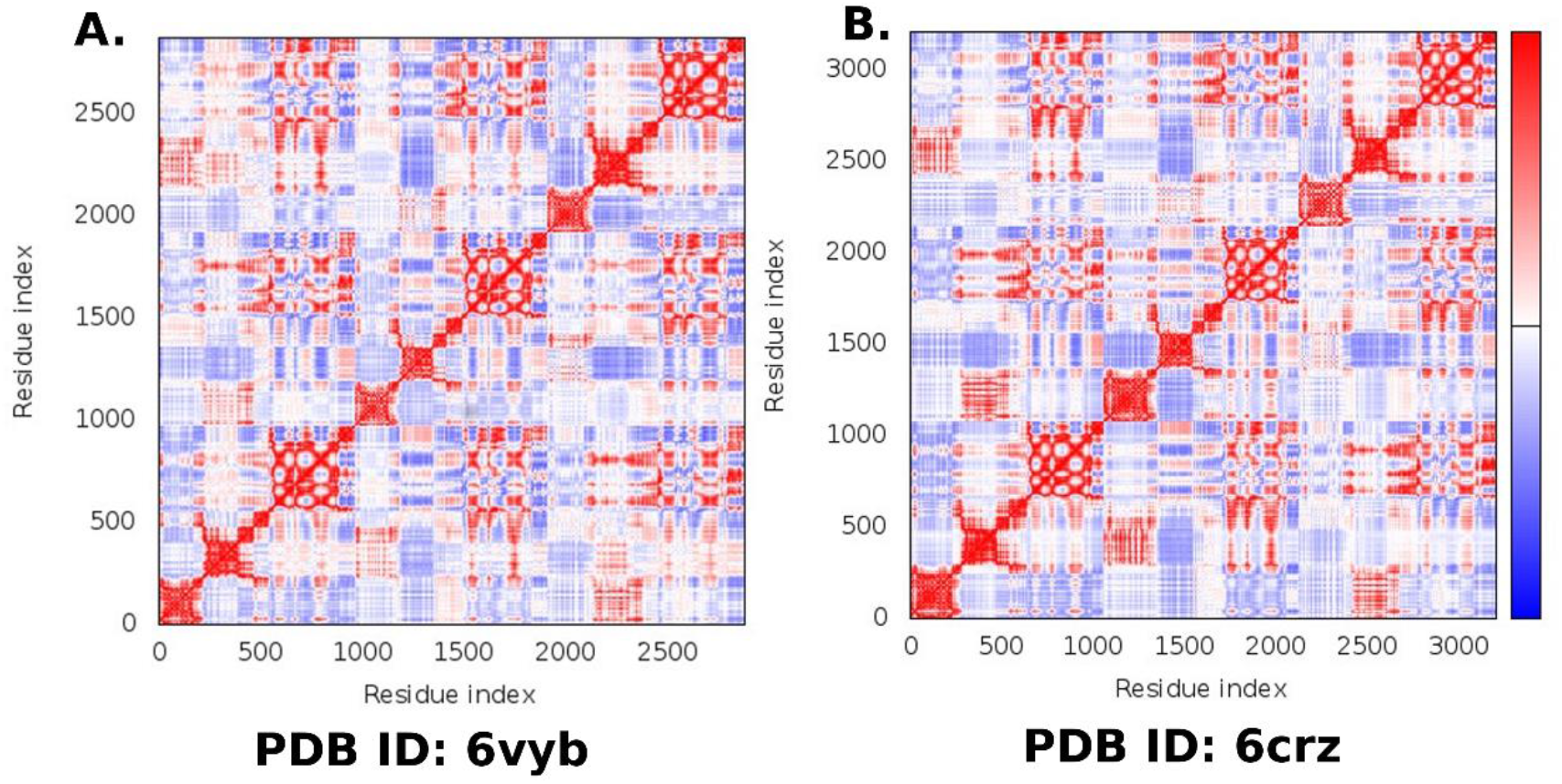
Coupling between pairs of residues-a covariance matrix with positively correlated (red), uncorrelated (white) or anti-correlated (blue) motions (prepared in iMODS webserver http://imods.chaconlab.org)

As usual, such differences are prominent in the low-frequency normal modes. To examine the internal residual coupling and its effects, we have studied low-frequency normal modes for RBD-up monomer of both the structures and noted that the second normal mode is capable to effectively capture these differences. It shows that RBD has relatively high amplitude motions for both the CoVs without hACE2 binding, while the S2-unit holds lower-amplitude motions (Fig. 4A). When we compare this prefusion up-form RBD local motion along the residues of CoV-1and CoV-2, it shows that the extended RBD region follows anti-phase like dynamics; CoV-2-RBD has positive eigenvectors in oppose to negative eigenvectors of CoV-1-RBD (Fig.4D). In contrast, these anti-phase dynamics are mostly vanished due to hACE2 binding. hACE2 binding makes the RBD relatively stable, with little faster movements of CoV-2-RBD (Fig.4A, C).

**Fig.4.**
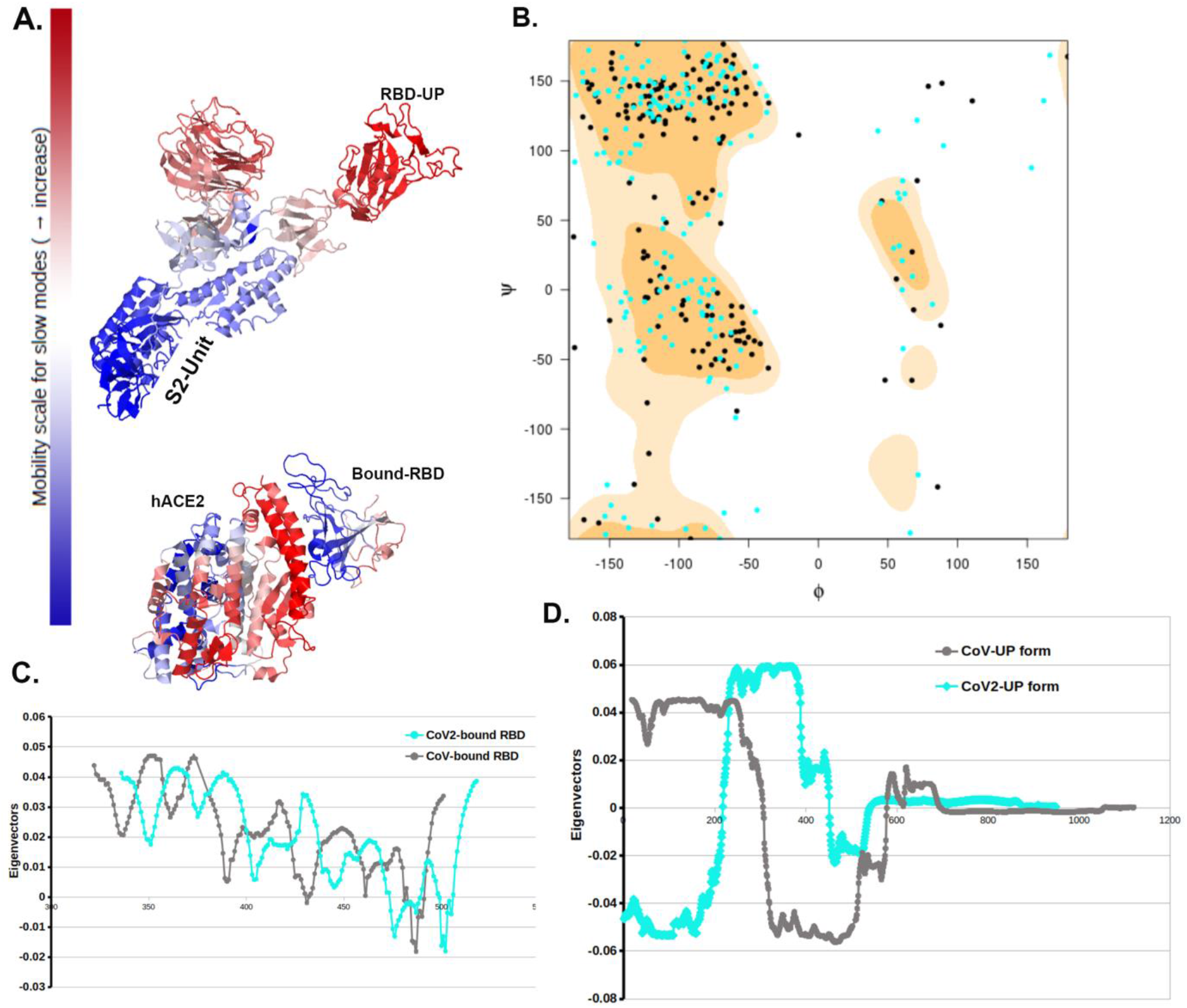
Normal Mode Analysis (NMA) and Ramachandran plot. We used low-frequency second normal mode to differentiate between SARS-CoV-1 and SARS-CoV-2 with and without hACE binding. (A) Its RBD-up conformation region in the S1-unit has higher mobility relative to other parts of the S-protein. In contrast, hACE2 binding make the RBD less mobile. (B) Ramachandran plot of AA residues of RBD bound with hACE2, shows significant conformational changes in the RBD. Cyan dots represent CoV-2-RBD AA residue phi-psi position, while black is for CoV-1complex. (C) Residue-wise eigenvectors of bound RBD AA residues is representing low-frequency motions. There are no significant differences in bound RBD motions between both the SARS-CoV-1 complexes. (D) Without hACE binding, RBD up-conformation protomer is showing significant differences in the domain motion in the RBD regions. It is like anti-phase-type dynamics between the two CoVs structures.

To further probe the conformational states and dynamical features of the whole spike protein and the impact of ACE2 binding, we applied a constrained geometric sampling method FRODAN [10, 42]. This method utilizes a coarse-grained molecular mechanics potential based on rigidity theory to explore receptor conformations well outside the starting structure. The method FRODAN (Framework Rigidity Optimized Dynamics Algorithm New) can be regarded as a low computational complexity alternative to MD simulations which can sample wide regions of high dimensional conformational space. We applied FRODAN on the whole spike protein with one RBD in the open up-state and the RBD-ACE2 complex of the single RBD domain bound to the ACE2.

The Figure 5A,B reveals RMSF fluctuations of the respective backbone atoms in both spike proteins at different energy cut-offs (temperatures). Consistent with the normal mode analysis, it is evident that for both spike proteins in SARS-CoV-1 and SARS-CoV-2, the RBD in the up conformation has higher conformational fluctuation relative to other domains. As we increase the hydrogen-bond energy cutoff (i.e. removing weak transient hydrogen bonds) conformational fluctuations in RBD tend to further increase. There is an overall increase in conformational dynamics in SARS-CoV-1 S structure in comparison to the SARS-CoV-2 (Table S2). In particular, RBD has higher RMSF in SARS-CoV-1. This is in agreement with rigidity analysis (Figure 8A, Table2), where we have shown that SARS-CoV-2 retains its overall rigidity better than SARS-CoV-1. Furthermore, large-scale computing efforts via Folding@home project have carried out millisecond MD simulation [40], where it was shown the RBD domain in the up state of the SARS-CoV-1 exhibits higher deviation from the respective crystal structure in comparison to the SARS-CoV-2 [40]. Interestingly, one of the RBDs in SARS-CoV-1 that is initially in the down configuration transitions to the open configuration. This correlates with previous studies which have shown that for the SARS-CoV-1, the two-up and three-up RBD states are also populated in the unbound spike protein [41]. On the other hand, for the SARS-CoV-2 previous studies indicated that the two-up and three-up states are rarely observed [43]. This trend is reflected also in our simulation results. For the SARS-CoV-2 the RBDs that are initially in the down state, remain in the closed state at wide range of energy cut-offs. Overall, it is evident that temperature increase affects the SARS-CoV-1 spike protein’s conformational stability in the more pronounced way than the SARS-CoV-2. This difference may provide clues why RBD binds tighter with ACE2 in SARS-CoV-2, and a general trend that the SARS-CoV-1 is more sensitive to environment conditions than the SARS-CoV-2 [12–14].

**Fig.5.**
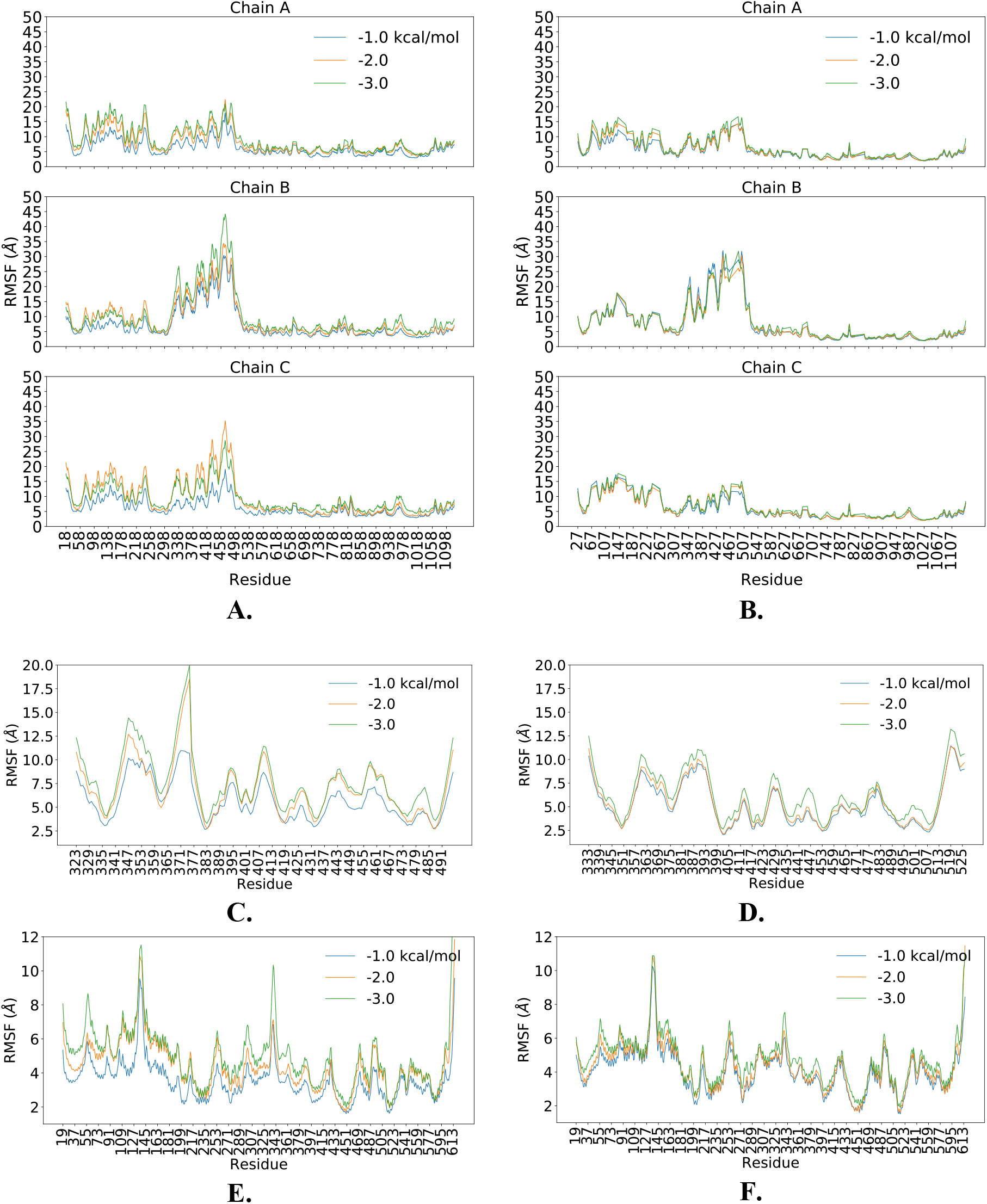
Conformational dynamics analysis using Monte-Carlo dynamics simulation method FRODAN [10]: (A, B) Backbone RMSF profiles of SARS-CoV-1 and SARS-CoV-2. The RBD domain that is initially in the up conformation is indicated by the filled area in pink. (C,D) RMSF of the RBD domain in the complex with ACE2. (E,F) RMSF of the ACE2 domain in the complex.

Figure 5C,D,E,F shows RMSF profiles of RBD and ACE2 in the complex at different energy cut-offs. Similarly, as in to the whole spike protein, the increase of the hydrogen-bond energy cutoff leads to the overall higher RMSF values in both complexes. The RBD domain fluctuates less in SARS-CoV-2. The average RMSF in SARS-CoV-2 is lower at all considered cut-offs (Table S3). This is supported by rigidity analysis findings (Figure 8B, Table2) that are indicating excessive flexibility in the SARS-CoV-1-ACE2 complex. Overall, it is evident that the stability of SARS-CoV-2-ACE2 complex results from the stronger interface contacts [14]. On the other hand, with the increase of the hydrogen-bond energy cutoff, ACE2 fluctuation magnitude increases in the similar manner in both SARS-CoV-1 and SARS-CoV-2 (Table S3).

### Uneven distribution of interface molecular interactions between the RBD and hACE2

Interface interactions certainly play crucial role for binding and that of dynamic domain separation shown in normal mode analysis. Here, we use Prodigy webserver to predict these interactions [15]. Interface between CoV-2-RBD and hACE2 is relatively higher than the SARS-CoV-1 complex, with higher potential for large intermolecular interactions. Although both complexes hold almost similar binding affinity for the hACE2, there are slight observable differences in dissociation coefficient (i.e. 7.3 nM for CoV-RBD-hACE2 complex and 12 nM for CoV-2-RBD-hACE2) (Table 1). This noted difference may arise due to differential interface molecular interactions (Table S1 and S2). In both complexes, almost all interface molecular interactions involve loops and small parts of beta-sheets secondary structures of RBD and a single alpha helical structure of hACE2, indicating interface instability (Fig.4). We find uneven distribution of interface interactions along this single hACE2 alpha-helix. While two end portions and the middle part of the helix hold most interactions, there are lot more differences of interacting residues between CoV-1and CoV-2 complex (Table S1). In particular, we note that hACE2.GLU35[OE2]–RBD.GLN493[NE2] and hACE2.THR27[O] – RBD.TYR489[OH] come closer (~2.6 Å) and form H-bonds in the CoV-2-RBD-hACE2 complex (Fig.6). Inclusion of these two additional polar interactions may affect subsequent proteolytic processing of S protein and membrane fusion with the S2-unit.

**Fig.6.**
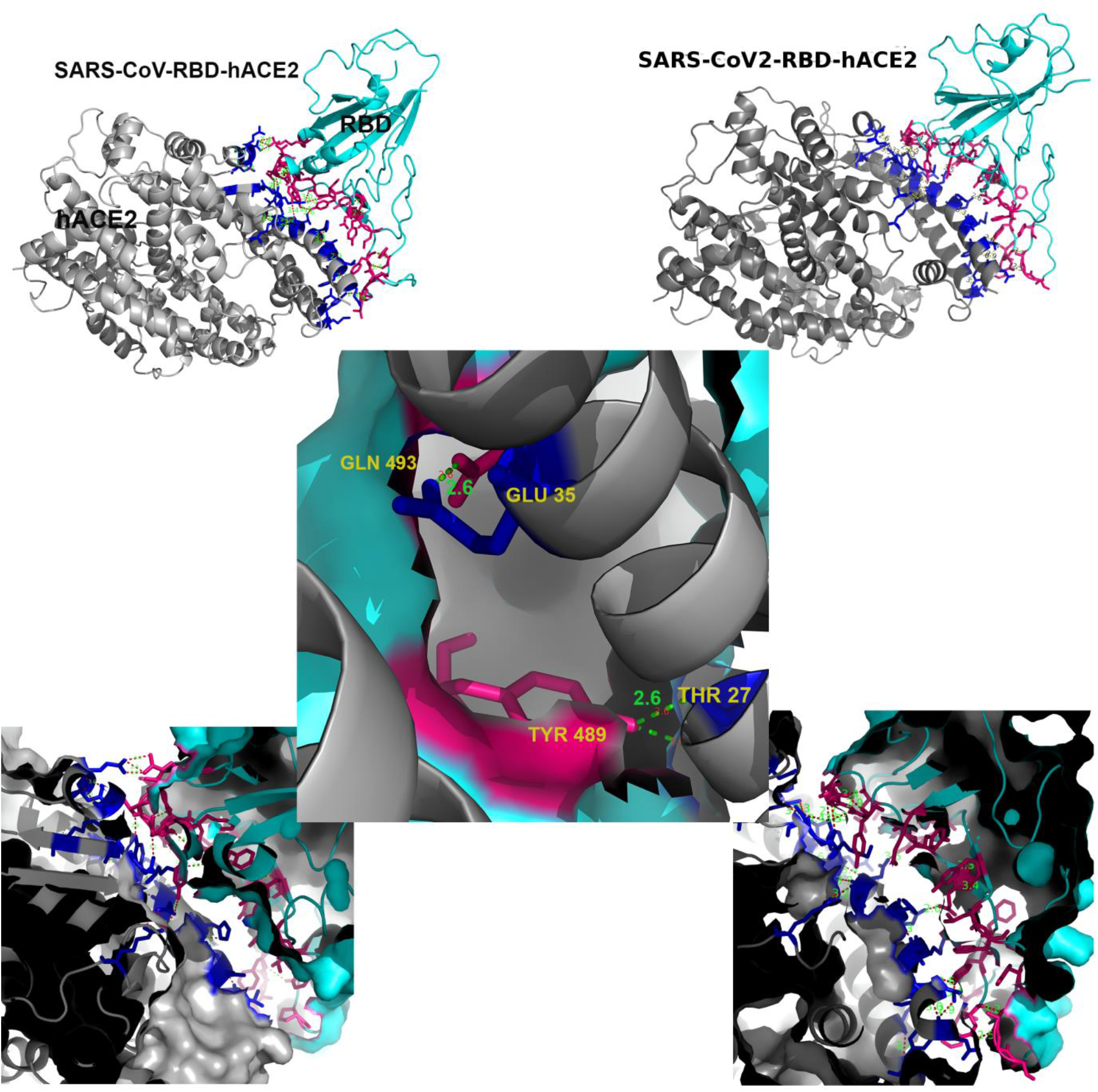
Interface interactions between CoV-1receptor-binding domain (RBD) and the human receptor membrane protein hACE2. In both the complexes, almost all interface molecular interactions involve loops and small parts of beta-sheets secondary structures of RBD and a single alpha helical structure of hACE2. We note that hACE2.GLU35–RBD.GLN493 and hACE2.THR27 – RBD.TYR489 (~2.6 Å) forms H-bond interactions in the CoV2-RBD-hACE2 complex.

**Table 1:** Interactions between hACE2 and the coronavirus Receptor-binding Domain (RBD)

| Complex name (PDB ids) | Bound-RBD surface area (Å<sup>2</sup>) | Interface Area (Å<sup>2</sup>) | Binding Affinity (kcal/mol) | Dissociation constant (nM) | Polar-polar contacts added | LJ interactions with water (kcal/mol) | Coulomb interactions with water (kcal/mol) |
| --- | --- | --- | --- | --- | --- | --- | --- |
| **SARS-CoV-RBD-hACE2** (2AJF) | 18435.672 | 796.7 | -11.1 | 7.3 | 2 | -281 | -4887 |
| **SARS-CoV2-RBD-hACE2** (6M17) | 19866.404 | 827.7 | -10.8 | 12 | 2+2; added<br>GLU35 – GLN493<br>THR27 – TYR489 | -261 | -5160 |

In order to check the reliability of the predicted interface interactions, we have calculated ^13^C chemical shifts (δ) using structural coordinates of both the CoV-1and CoV-2 hACE2 bound complex. This calculation is carried out using SHIFTX2 that combines ensemble machine learning approaches with the sequence alignment-based methods and that has carefully been trained with 197 high-resolution protein structures. We note that there is no overall significant differences in δ (ppm) between the hACE2 bound CoV-1and CoV-2 complex (Fig.7A). However, relative absolute differences in δ (Δδ) are prominent at the interface between hACE2 and RBD (Fig.7A middle). Detected uneven distribution and changes of interface interactions are shown to be strongly correlated to deviation in the chemical shifts (Δδ). In particular, the hACE2 interface helix and the interacting RBD domain of the CoV-2 RBD shows significant difference in Δδ (Fig.7B). As expected, very high Δδ values are noted at and nearby H-bond forming residues GLU 35, THR 27 on the hACE2 helices and GLN 493 and TYR 489 on the RBD interface (Fig.7B). A large Δδ pick of 14.86 ppm on the RBD-interface and 2.50 ppm on the hACE2 helix are noted, with significant changes in the middle and end parts of the interacting interface helix. Furthermore, large picks in Δδ corresponds to AA substitutions. For example, Δδ pick of 14.86 and 8.90 ppm correspond to the substitution THR433GLY446 and THR487ASN501 on the interacting RBD domain.

**Fig.7.**
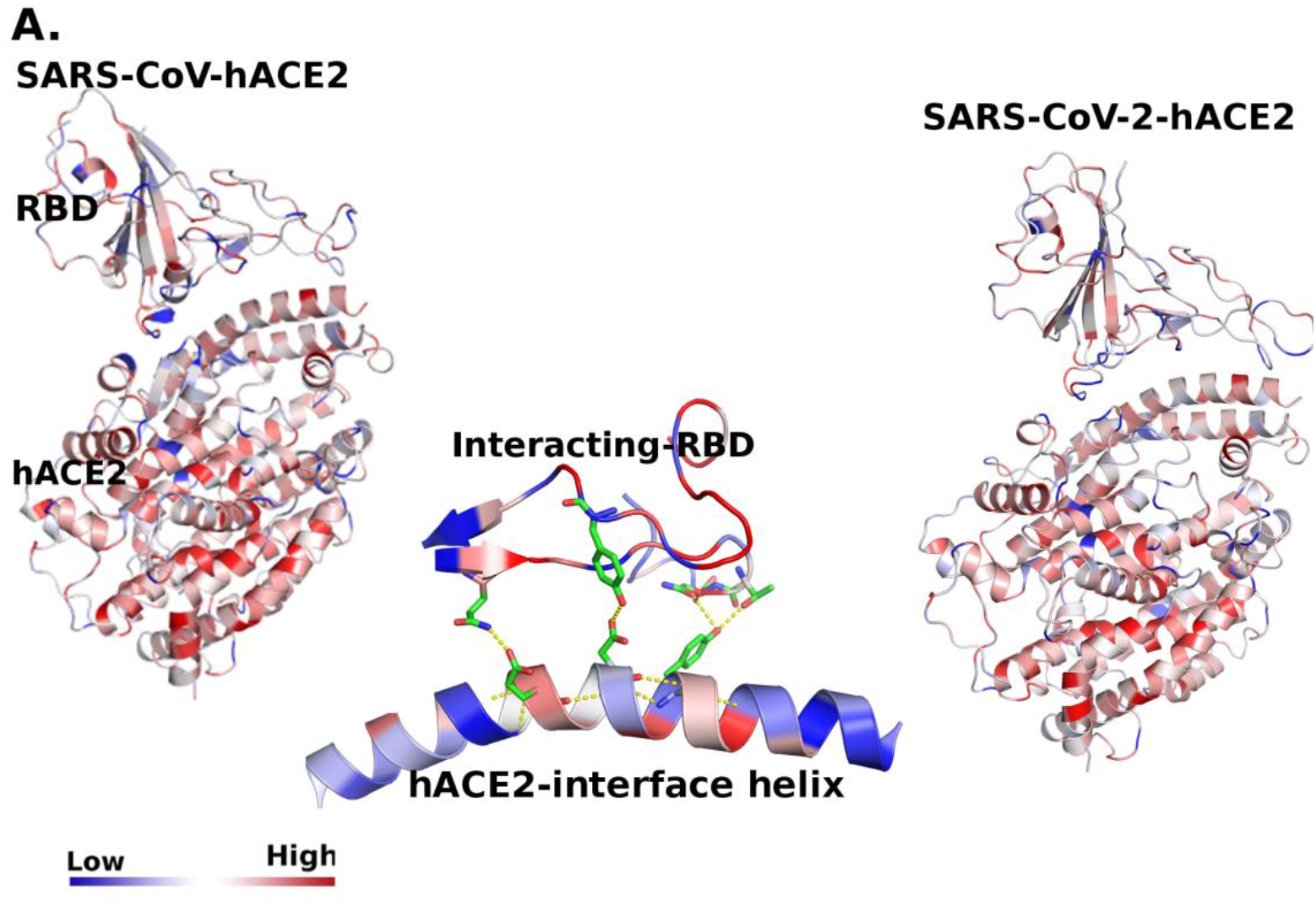
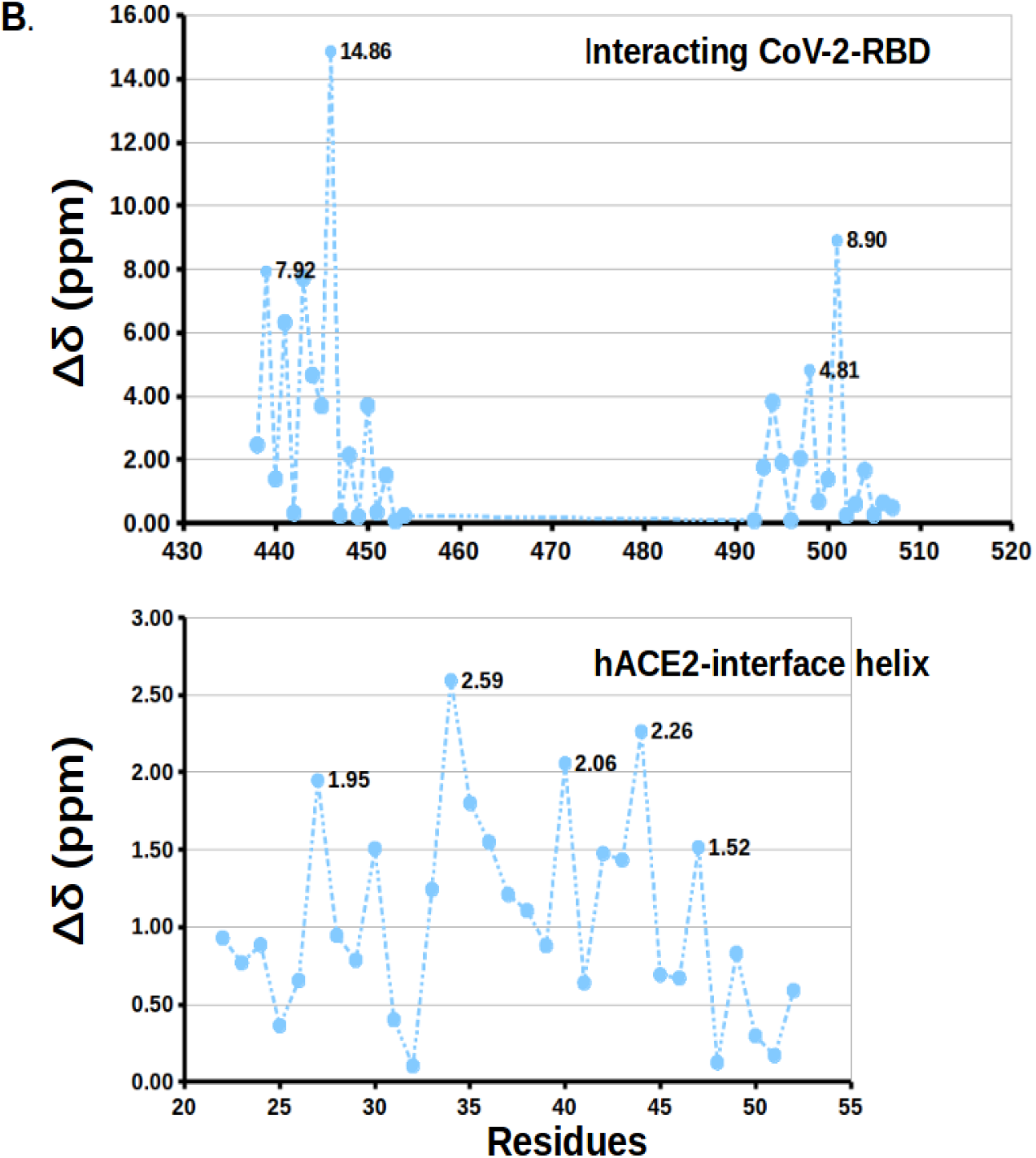
^13^C Chemical shifts (δ) and changes in ^13^C chemical shifts (Δδ) indicate the changes in the interface interactions between hACE2 and RBD. (A) there are no overall significant differences in δ (ppm) between the hACE2 bound CoV-1and CoV-2 complex (top two figures), the middle figure shows the deviation in chemical shifts (Δδ) at the interface interacting sites, (B) A large change in Δδ are observed at and nearby H-bond forming residues GLU 35, THR 27 on the hACE2 helices and GLN 493 and TYR 489 on the RBD interface.

### hACE2 binding induces changes in structural flexibilities

Structural flexibility of the spike indicates its potential to bind and interact with the human receptor protein hACE2. To probe this we have carried out rigidity analysis using Floppy Inclusion and Rigid Substructure Topography (FIRST) program which is based on pebble game algorithm and techniques in rigidity theory. In this approach, we first create a constraint network, where the spike protein is modelled in terms of nodes (atoms) and edges (covalent bonds, hydrogen bonds, hydrophobics etc). A hydrogen bond cutoff energy value is selected where all bonds weaker than this cutoff are ignored. The strength of hydrogen bonds is calculated using a Mayo energy potential. The spike protein network is next decomposed into rigid clusters and flexible regions. We used FIRST to decompose the spike protein structures into rigid clusters with the input of H-bond energy cutoff (H-cut-off, kcal/mol). With H-cut-off of −0.1 −0.5, −1.0, −1.5, −2.0, −2.5, and −3.0, CoV-1 and CoV-2 spike with- and without hACE2 bound have been decomposed. Number of rigid clusters and degree of freedoms consistently increase with increasing H-cutoff. Without ACE2 bound, CoV-1 S have been decomposed into higher number of rigid clusters with much higher degrees of freedom relative to CoV-2 S (Figure 8 and Table 2). This strongly indicates that CoV-1 S become relatively much flexible, generating a large conformational ensemble with less potential for binding with hACE2. This result is consistent with the recent observations in an extensive 0.1 second MD simulation which noted a large opening of spike with presence multiple cryptic epitopes [40]. In contrast, when RBD binds with hACE2, CoV2-RBD-hACE2 becomes relatively more rigid than CoV1-RBD-hACE2, which favours proteolytic processing for membrane fusions.

**Fig.8.**
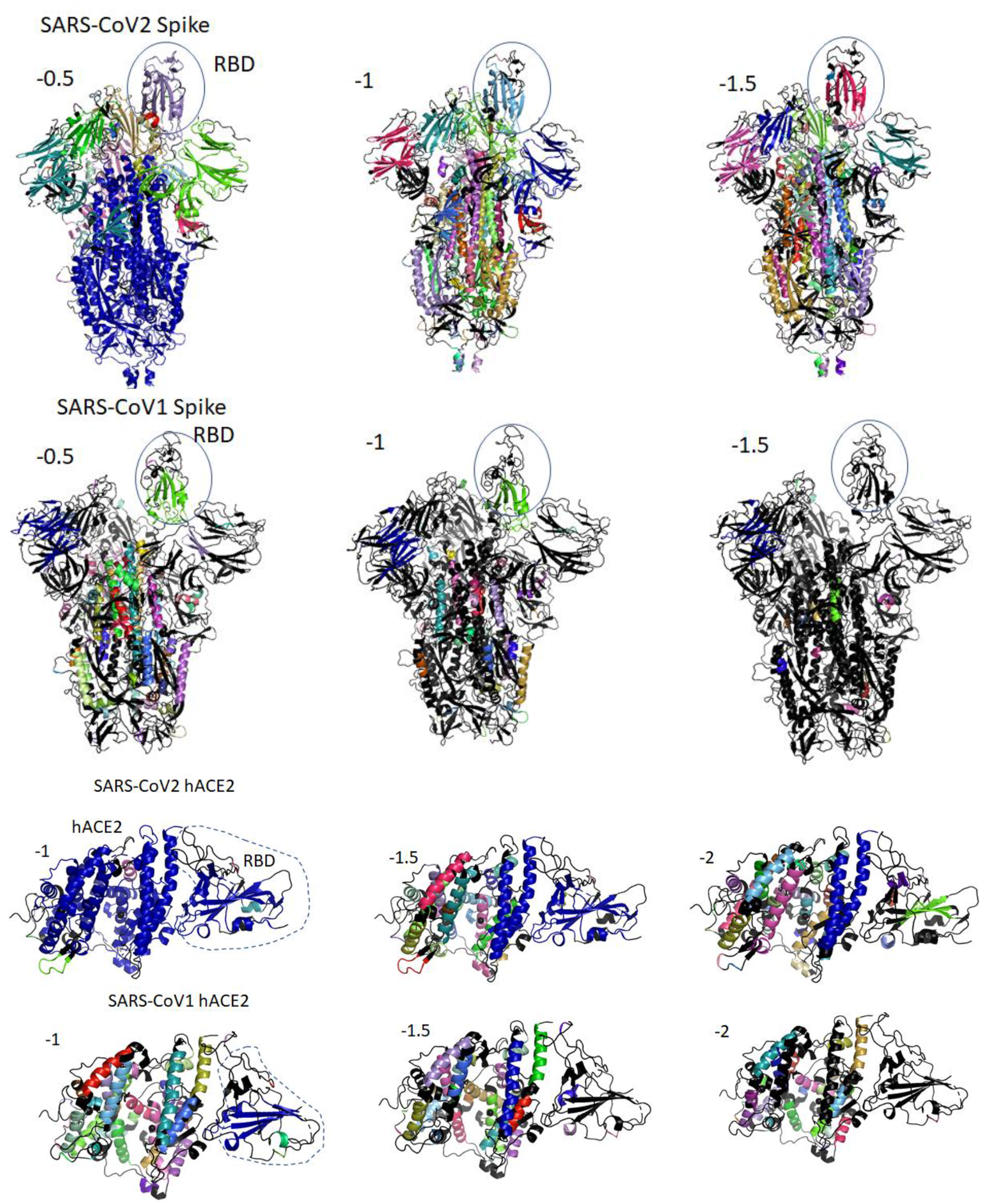
Rigidity analysis of protein structures. Rigid cluster decomposition analysis using rigidity-theory approach FIRST on Spike protein in SARS-CoV1 vs SARS-CoV2. Individual colours indicate presence of large rigid clusters (blue being the largest rigid cluster), while black colour represents flexible regions. At a low energy cutoff (−0.5 kcal/mol) SARS-CoV2 is significantly more rigid than SARS-CoV1, as it is dominated by a few large rigid clusters. As we increase the energy cutoff, SARS-CoV1 continues to gain more flexibility while SARS-CoV2 retains most of its rigidity. At −1.5 kcal/mol SARS-CoV1, including its RBD, has lost almost all internal structural rigidity, while SARS-CoV2 still maintains significant rigidity. Rigid cluster decomposition analysis using rigidity-theory approach FIRST on SARS-CoV1 vs SARS-CoV2 RBD-hACE2 complex. SARS-CoV2 RBD-hACE2 complex at −1kcal/mol is dominated by one large rigid cluster, whereas SARS-CoV1 RBD-hACE2 complex is more flexible consisting of several rigid cluster. As hydrogen bond energy cutoff is increased, SARS-CoV1 RBD-hACE2 complex is losing its rigidity more rapidly than SARS-CoV2 RBD-hACE2 complex.

**Table 2:**
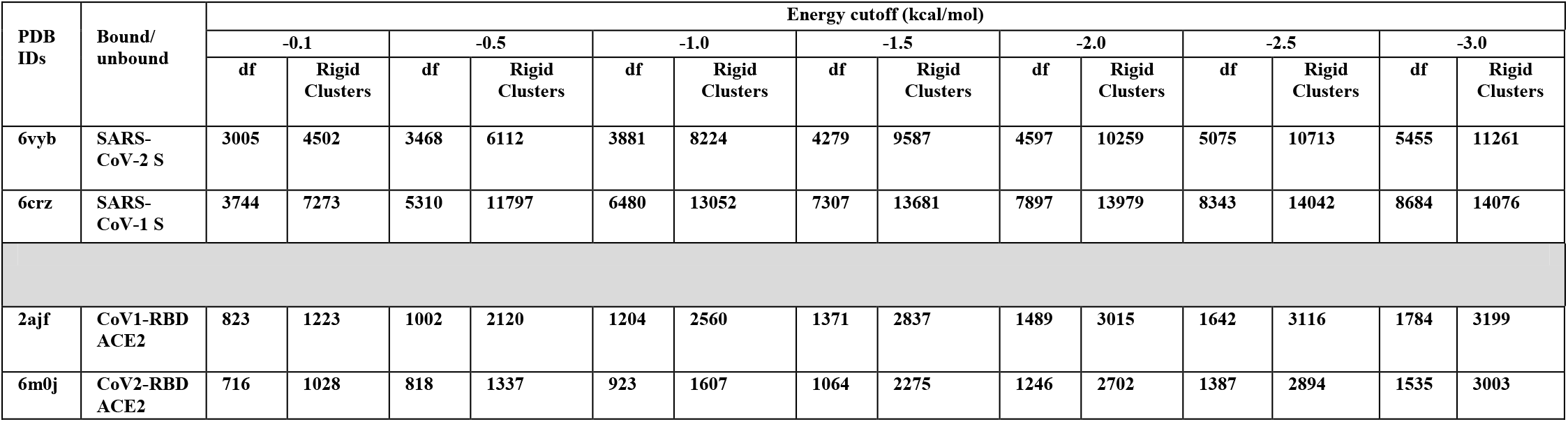
Summary of rigidity analysis

### Interfacial water promotes and stabilizes the formation of CoV-2/hACE2 complex

Previous studies [17,18] have indicated that water-mediated interactions can play an essential role in stabilization of CoV-1/hACE2 complex. It can be shown [18] that a gap between the RBD/hACE2 protein-protein complex includes several hundred interfacial water molecules which can form bridges between amino acids of the RBD and hACE2 subunits. Using the three-dimensional reference interaction model (3D-RISM) approach (see details in Methods) we have evaluated this phenomenon and calculated a distribution of interfacial water and its probable influence on the complex formation. Figure 9a demonstrates a water distribution in the interfacial region for CoV-2/hACE2 complex. Water molecules are depicted only in the most probable positions. As it is seen, the interfacial water interacts strongly with the RBD as well as with the hACE2 subunits, the latter confirms the previous molecular dynamic simulations [18]. However, our calculations yield the water-mediated interactions to be stronger for the CoV-2/hACE2 complex than those for the CoV-1/hACE2. See for example Fig. 9b demonstrating a relative difference in distributions of water oxygens (blue) and hydrogens (red) between the two complexes. Moreover, the water interacts more strongly for CoV-2/hACE2 rather than for pure coronavirus. This effect is also confirmed by the calculations of water-complex interaction energies (Table 1). The data on interaction energies between the complexes and interfacial water proves this fact. Although the Lennard-Jones energy between the CoV-1/hACE2 and water is stronger than that for the CoV-2/hACE2 complex, the total interaction energy has a more profit for the latter complex due to a strong decrease in the electrostatic energy. This observed effect can be explained by a polarization of interfacial water which is stronger for the CoV-2/hACE2 complex.

**Fig.9.**
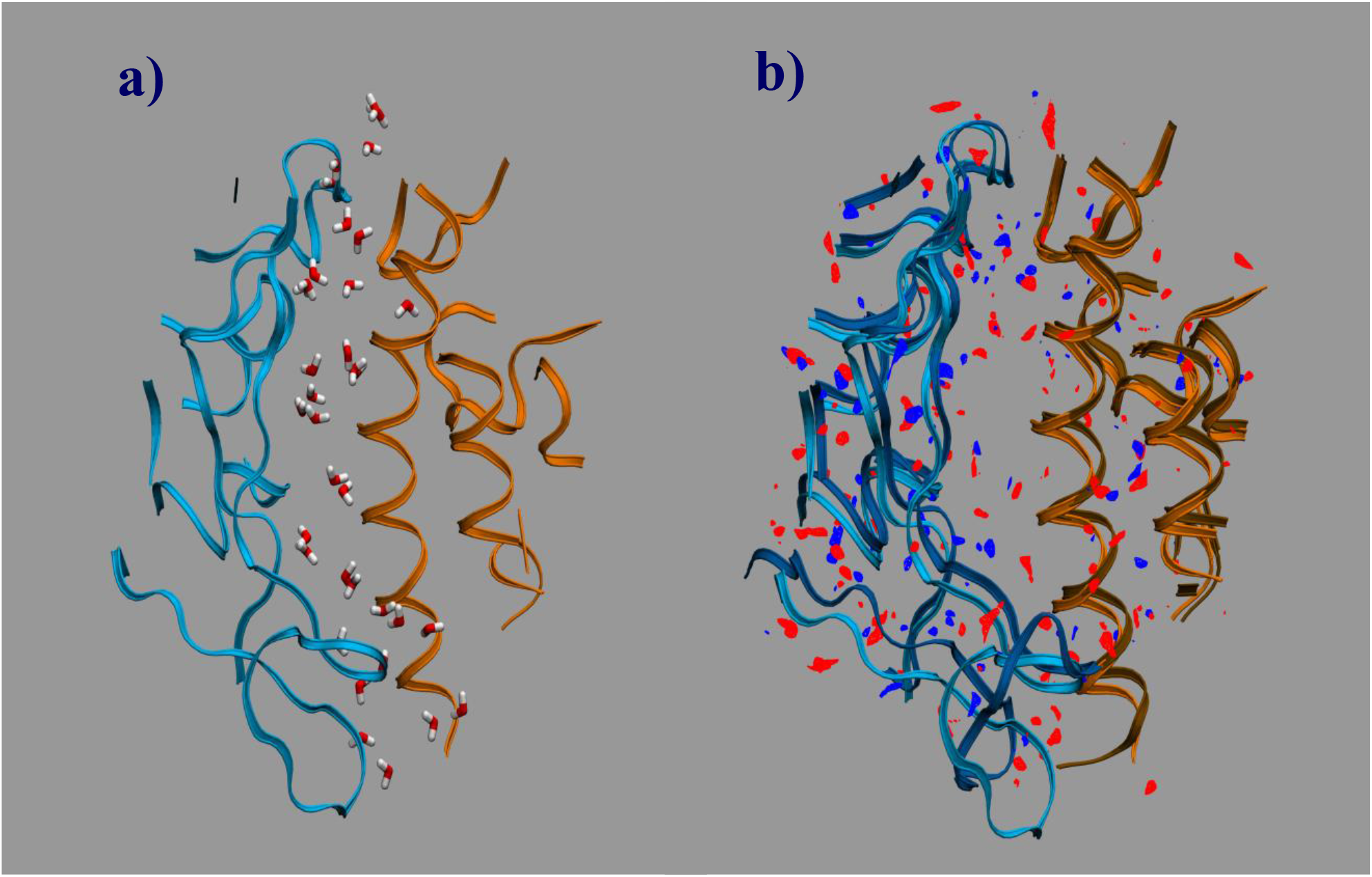
Water distribution in the interfacial region of complexes: a) for the CoV-2/hACE2 complex, b) differences in distributions of water oxygens (blue colour) and water hydrogens (red colour) between the CoV-2/hACE2 and CoV/hACE2 complexes. The RBD is indicated by blue ribbons, the hACE2 by orange ribbons, and the CoV/hACE2 is shown as background for the differences.

Apart from it, the latter complex forms water bridges whose number is larger by 2 than in the case of CoV-1/hACE2. At the same time the water bridges is more stronger in the CoV-2/hACE2 aggregate. For example, we have calculated the potential of mean force (pmf) between atoms of corona virus and hACE2 receptor (Fig. 10). It is clearly seen that bridging water molecule forms simultaneously hydrogen bonds with CoV-2 (Gln493) and hACE2 (Glu35) amino acids in the case of CoV-2 (Figs. 10a and 10c). However, the similar hydrogen bond with bridging water is very weak and diffusive for the CoV-1/hACE2 complex (Figs. 10b and 10d). Therefore, we conclude that bridge water molecules play a significant and, perhaps, crucial role in stabilizing CoV-2/hACE2 complex.

**Fig.10.**
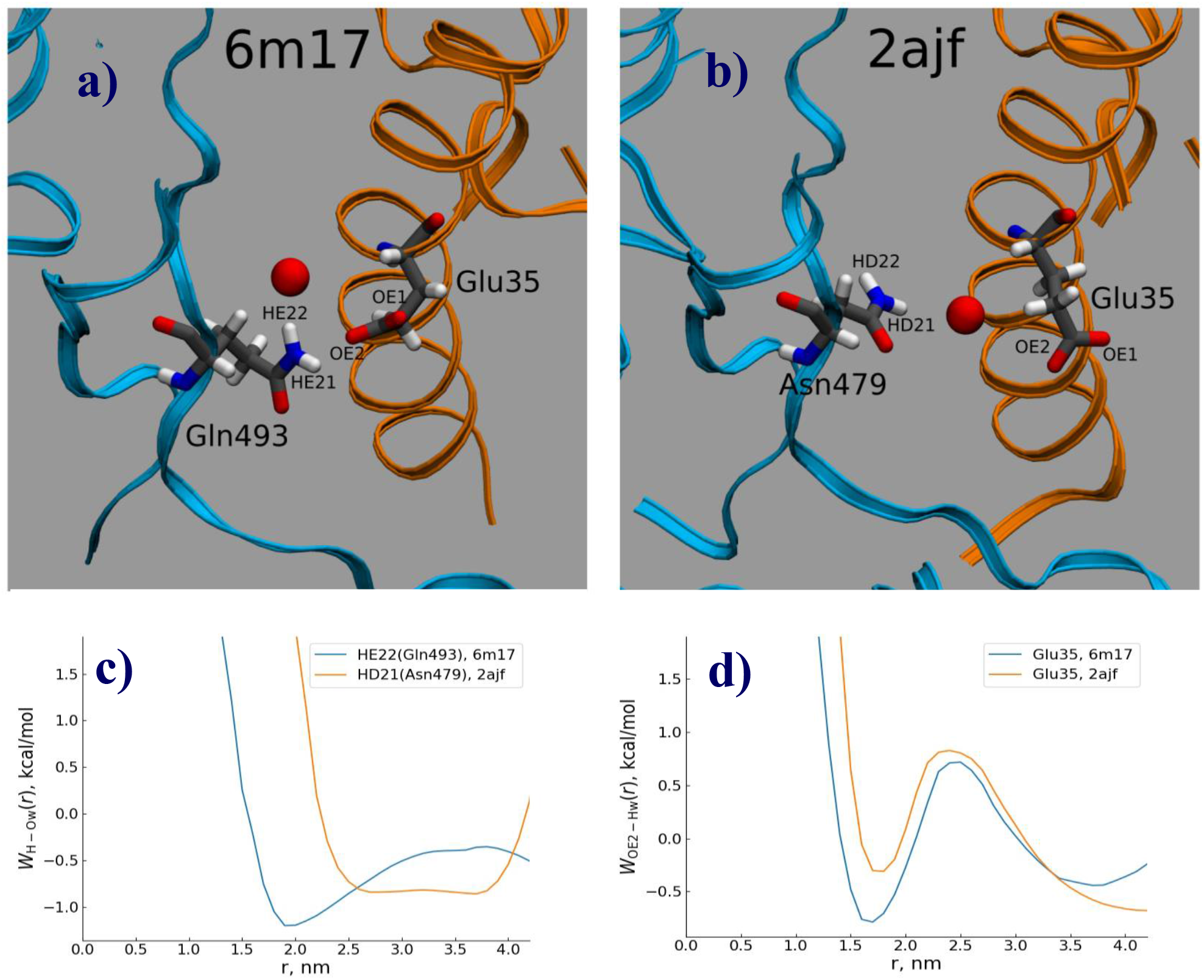
Water bridging the complexes: a) location of water oxygen bridging the CoV-2/hACE2 complex, b) the same for the CoV-1/hACE2 complex, c) the pmf of H-O distribution for the CoV-2 and CoV-water bridges, and d) the pmf of O-H for the hACE2-water bridges in the complexes.

## Conclusions

Viral entry into host cell, subsequent replication and transmissibility are essentially dependent on receptor recognition, binding, and membrane fusion mechanisms. SARS-coronavirus use ectodomain trimer, spike glycoprotein (S-protein), to mediate this viral entry [19]. Both SARS-CoV-1and SARS-CoV-2 recognizes the human membrane receptor protein hACE2 through its receptor-binding domain (RBD) positioned within the flexible S1 unit of S-protein protomer. S-protomer with the RBD up-conformational forms can only recognize hACE2. With down RBD conformations in all the three protomers of S-protein prevent hACE2 binding. Prefusion RBD conformation is, therefore, very important to determine receptor binding and thereby its differences between CoV-1and CoV-2 underpin the higher success of human-to-human transmission of SARS-CoV-2. Understanding prefusion conformational dynamics as well as its binding mechanism to the receptor among closely related species is critical for designing vaccine and inhibitors to strop viral entry. For example, one recent structural study has tested multiple SARS-CoV-1RBD specific monoclonal antibodies and found that none of them have notable binding to CoV-2 RBD [5]. Recent major therapeutic efforts are targeting the interactions between the SARS-CoV-2 spike (S) glycoprotein and the human cellular receptor angiotensin-converting enzyme, hACE2 [20–22]. It has recently been demonstrated that SARS-CoV-1S murine polyclonal antibodies can potentially inhibit SARS-CoV2 S mediated viral entry. However, its underlying inhibition mechanism is poorly understood [4].

Using recently reported cryo-EM structures of SARS-CoV-1 and CoV-2 S protein we find effective conformational differences in the RBDs at prefusion state. The RBD-up conformation in the SARS-CoV-1 is less stable than in the SARS-CoV-2 which is indicated by the excessive flexibility in the whole structure. When bound with higher surface area of CoV-2 RBD and large local conformational changes in the AA residues, it establishes higher interactions with the human receptor hACE2. These underlying conformational differences are illustrated with the higher RMSD of Cα atoms, changes in phi and psi dihedral angles and relatively large amplitude anti-phase-like RBD motion. In particular, we note that hACE2.GLU35– RBD.GLN493 and hACE2.THR27 – RBD.TYR489 come closer in the CoV2-RBD-hACE2 complex and forms important H-bonds. However, it remains to be explored how efficiently CoV-2 regulate and open-up RBD for hACE2-binding. The RBD of CoV-2 forms stronger water bridges with the hACE2, and it plays a significant role in the total stabilization of the CoV-2/hACE2 complex. Inclusion of the two noted additional polar interactions affects structural flexibilities. We find such interaction changes critically affect structural flexibility. In particular, hACE2 binding makes the CoV-2 structure more rigid relative to CoV-1complex. This h-ACE2-induced changes in structural flexibility favours subsequent proteolytic processing which is essential for membrane fusion. This understanding certainly provides a clue for designing novel vaccine and antiviral drugs.

## Methods

### Root-mean-squared deviation (RMSD) of Cα atoms

We used MatchMaker in UCSF-Chimera to superimpose structures. It performs a best fit after automatically identifying pair of residues. It considers similar structures to be superimposed while there are large sequence dissimilarities by producing residue pairing uses with the inputs of AA sequences and secondary structural elements. After superposition, it uses the usual Euclidian distance formula for calculating RMSD in angstrom unit with the use of PDB atomic coordinates files.

### Normal Mode Analysis (NMA)

DynOmics ENM Server [23] was used to perform normal mode analysis (NMA). NMA is a well-explored technique for exploring functional motions of proteins. It combines two elastic network models (ENMs) — the Gaussian Network Model (GNM) and the Anisotropic Network Model (ANM) to evaluate the dynamics of structurally resolved systems, from individual molecules to large complexes and assemblies, in the context of their physiological environment. In the GNM model, network nodes are the C-alpha atoms and the elastic springs represented the interactions. We used GNM with interaction cut-off distance of 7.3 Å and spring constant scaling factor cut-off of 1 Å for the calculation of the elastic network model. We calculated the first twenty slowest modes for all the CoVs structures. The eigenvectors of these modes represent the global motions and the constrained residues help in identifying critical regions such as hinge-bending regions and thereby giving an idea of domain motions around these regions. We plotted the second slowest mode in different conditions which showed a significant difference in motions.

### Constrained geometric simulations for conformational sampling

To study dynamics of the considered structures, we employed a low computational complexity alternative to MD geometric molecular simulation package FRODAN v1.0.1 [10]. We run FRODAN in the non-targeted mode, generating 30,000 candidate structures for each case. The temperature impact was evaluated by running simulations at different hydrogen bond energy cut-offs. The considered range of the energy cut-offs was from −1.0 to −3.0 kcal/mol, with that of the more negative cut-off values correspond to higher temperature. During each individual run the cut-off value was kept constant. Before running simulations, hydrogen bonds to each considered structure were added using MolProbity server: (http://molprobity.biochem.duke.edu/).

### Rigidity analysis

We use the program Floppy Inclusion and Rigid Substructure Topography (FIRST) [16] that generates a constraints network from the coordinates of the protein structure in terms of nodes (atoms) and edges (covalent bonds, hydrogen bonds and hydrophobic interactions). Hydrogen bonds were ranked according to their energy strength using Mayo potential, whereas the value of energy strength was selected in such way that the bonds strength below this cut off were ignored. First then applies the rigorous mathematical theory of rigid and flexible molecular structure and pebble game algorithm calculates the degree of freedom of motion to rapidly decompose a protein into rigid clusters and flexible region.

### 3D RISM calculations

The 3D-RISM method [24–25] with the 3D-Kovalenko-Hirata closure was used to describe hydration of CoV-2/hACE2 and CoV/hACE2 complexes. These calculations describe hydration on a molecular-atom level and has proven their power in many successful studies of ion / molecule hydration, including various biocompounds [26–36]. In the framework of 3D-RISM theory molecule-atom spatial distribution functions are obtained. These functions are the 3D-site distribution functions of water atoms around the complex. The RISM-calculations were carried out for the studied complexes hydrated in water at ambient conditions. For water the modified version of the SPC/E model (MSPC/E) was used [37]. The corresponding LJ parameters of the solute atoms were taken from the ff14SB force fields [38]. The 3D-RISM calculations were performed using the rism3d.snglpnt codes from the AmberTools 16 package [39]. The 3D-RISM equations were solved on a 3D-grid of 350 × 320 × 350 points with a spacing of 0.025 nm. A residual tolerance of 10^−6^ was chosen. These parameters are enough to accommodate the complex together with sufficient solvation space around it so that the obtained results are without significant numerical errors.

## Acknowledgments

Our thanks to our lab members, Sikkim University for their support while completing a part of computational study in this work. The study of ChGN was supported by the Government Contract of the Institute of Theoretical and Experimental Biophysics, Russian Academy of Sciences (N 075-00381-21-00). The study of MVF and SEK was performed under the Government Contract of G.A. Krestov Institute of Solution Chemistry, Russian Academy of Sciences (N 0077-2019-0033).

## Author Contributions

AC and AS have conceived the study. NK, SB, MR, AC have conducted computational experiments and analytical analysis. AS, KS, AT carried out rigidity theory and Monte Carlo simulation analysis based on PDB files and has written the corresponding part. GC, MF, SEK has formulated the problem of hydration for COVID-hACE2 complex, performed the initial calculations and written the corresponding section of the paper. SEK has performed all 3D-RISM calculations and a visualization of the results. MVF has conducted a formal analysis and discussion of 3D-RISM results and written the corresponding section of the paper. GC, AS, and AC have reviewed and written the article.

## Supplementary Tables and Figures

**Table S1:** Interface residual interactions between CoV-1S glycoprotein receptor-binding domain (RBD) and human ACE2.

SARS-CoV-RBD-hACE2 Complex
|  | Human Receptor | Virus RBD Domain | Affinity (PRODIGY) |
| --- | --- | --- | --- |
| 1 | TYR83 | ASN473 | The binding affinity ($\Delta G$) is $\Delta G = -11.1$ (kcal mol<sup>-1</sup>) |
| 2 | GLN24 | ASN473 |  |
| 3 | ASP38 | TYR436 |  |
| 4 | GLU37 | TYR491 |  |
| 5 | LYS353 | TYR481 |  |
| 6 | LYS353 | GLY482 | The dissociation constant ($K_d$) at 25.0 °C is:<br>$K_d = 7.3E-09$ (M)<br>= 7.3 nM |
| 7 | LYS353 | GLY488 |  |
| 8 | TYR41 | THR486 |  |
| 9 | GLU329 | ARG426 |  |

|  | Human Receptor | Virus RBD Domain | Affinity (PRODIGY) |
| --- | --- | --- | --- |
| 1 | LYS353 | GLY502 | The binding affinity ($\Delta G$) is $\Delta G = -10.8$ (kcal mol<sup>-1</sup>) |
| 2 | TYR41 | THR500 |  |
| 3 | TYR41 | ASN501 |  |
| 4 | ASP38 | TYR449 |  |
| 5 | GLU35 | GLN493 |  |
| 6 | THR27 | TYR489 | The dissociation constant ($K_d$) at 25.0 °C is:<br>$K_d = 1.2E-08$ (M)<br>= 12 nM |

**Table S2:**
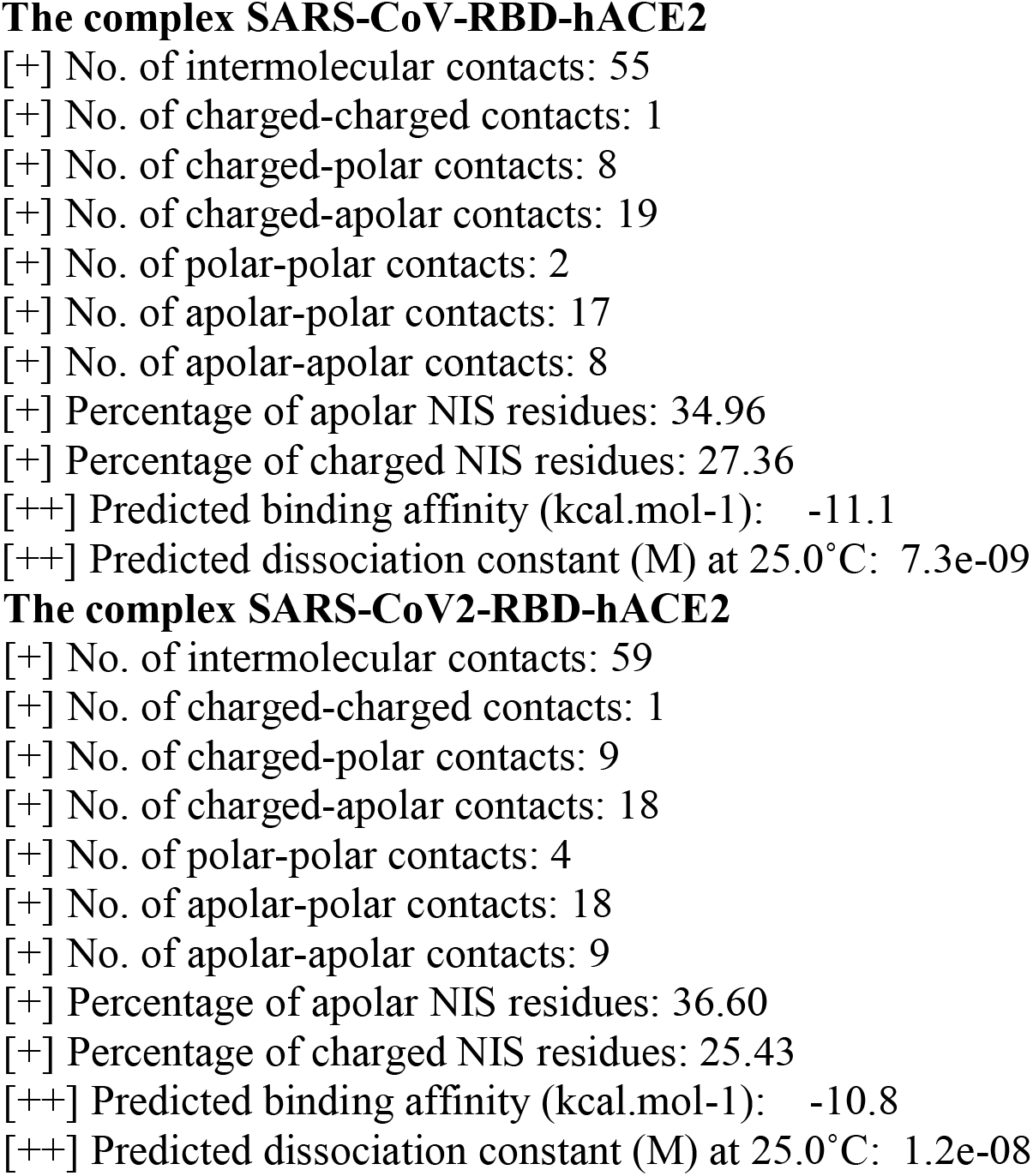
Interface interaction summary between Receoptor-binding domain (RBD) of CoV-1and human ACE2.

**Table S2.** Average RMSF values of spike protein with RBD in the up state for SARS-CoV-1 and SARS-CoV-2. Highest value among SARS-CoV-1 and SARS-CoV-2 at selected energy cut-off is indicated in bold.

|  | All domains |  | RBD up |  |
| --- | --- | --- | --- | --- |
| Cut-off | SARS-CoV-1 | SARS-CoV-2 | SARS-CoV-1 | SARS-CoV-2 |
| -1.0kcal/mol | **6.5±3.9Å** | 5.9±4.7Å | 17.7±5.6Å | **18.0±7.9Å** |
| -2.0kcal/mol | **8.7±5.6Å** | 6.1±4.3Å | **20.1±6.2Å** | 16.4±6.8Å |
| -3.0kcal/mol | **9.4±5.8Å** | 6.5±4.4Å | **24.8±7.9Å** | 16.6±6.8Å |

**Table S3.**
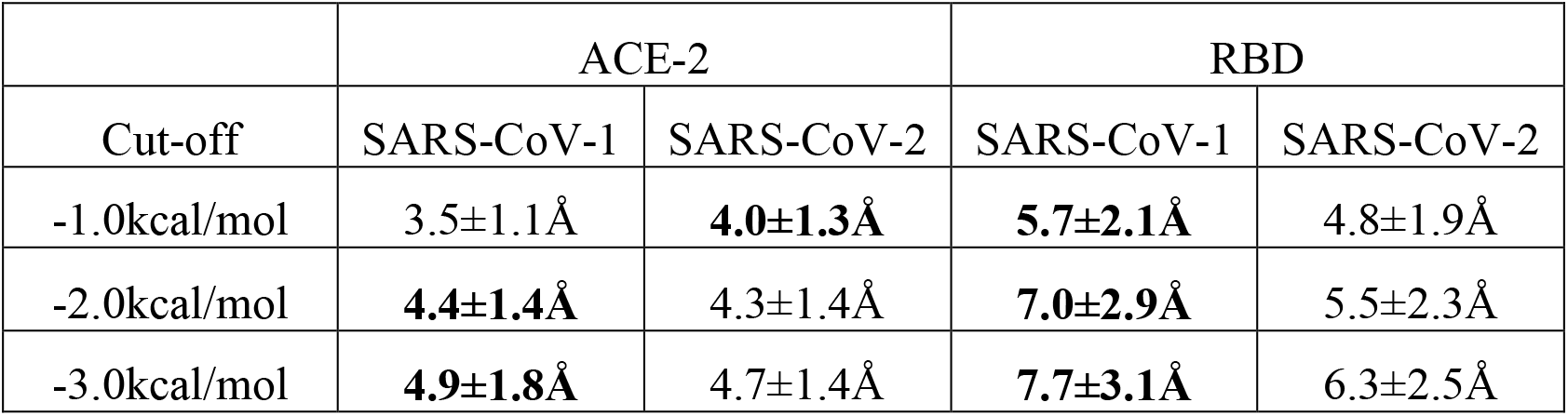
Average RMSF values of the ACE2-RBD complex of SARS-CoV-1 and SARS-CoV-2. Highest value among SARS-CoV-1 and SARS-CoV-2 at selected energy cut-off is indicated in bold.

|  | ACE-2 |  | RBD |  |
| --- | --- | --- | --- | --- |
| Cut-off | SARS-CoV-1 | SARS-CoV-2 | SARS-CoV-1 | SARS-CoV-2 |
| -1.0kcal/mol | 3.5±1.1Å | **4.0±1.3Å** | **5.7±2.1Å** | 4.8±1.9Å |
| -2.0kcal/mol | **4.4±1.4Å** | 4.3±1.4Å | **7.0±2.9Å** | 5.5±2.3Å |
| -3.0kcal/mol | **4.9±1.8Å** | 4.7±1.4Å | **7.7±3.1Å** | 6.3±2.5Å |

**Fig.S2.**
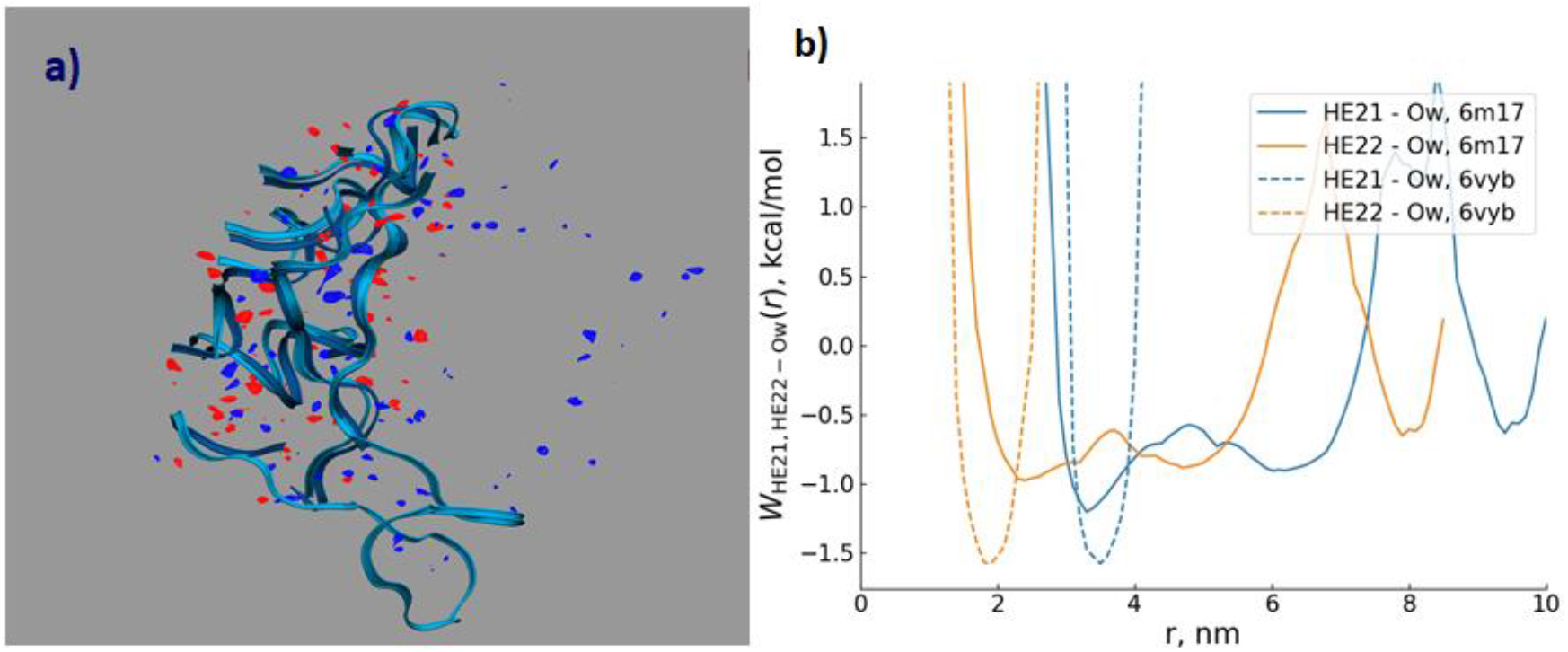
Water distribution around RBD of SARS-CoV-2. (a) Differences in distributions of water oxygen (blue) and water hydrogens (red) between the CoV-2/hACE2 and CoV-2; (b) the pmf RBD-O and RBD-H for the CoV-2 and CoV-2/hACE2 complex. The RBD is indicated by blue ribbons, the RBD of CoV-2 is shown as background for the differences.

